# High-throughput Single-cell CNV Detection Reveals Clonal Evolution During Hepatocellular Carcinoma Recurrence

**DOI:** 10.1101/2020.12.09.417626

**Authors:** Liang Wu, Yuzhou Wang, Miaomiao Jiang, Biaofeng Zhou, Yunfan Sun, Kaiqian Zhou, Jiarui Xie, Yu Zhong, Zhikun Zhao, Michael Dean, Yong Hou, Shiping Liu

## Abstract

Single-cell genomics provides substantial resources for dissecting cellular heterogeneity and cancer evolution, but classical DNA amplification-based methods are low-throughput and introduce coverage bias during sample preamplification. We developed a **s**ingle-**c**ell **D**NA library preparation method without **p**reamplification in **n**anolitre scale (scDPN). The method has a throughput of up to 1,800 cells per run for copy number variation (CNV) detection. Also, it has a lower level of amplification bias and noise than the multiple displacement amplification (MDA) method and showed high sensitivity and accuracy based on evaluation in cell lines and tumour tissues. We used this approach to profile the tumour clones in paired primary and relapsed tumour samples of hepatocellular carcinoma (HCC). We identified 3 clonal subpopulations with a multitude of aneuploid alterations across the genome. Furthermore, we observed that a minor clone of the primary tumour containing additional alterations in chromosomes 1q, 10q, and 14q developed into the dominant clone in the recurrent tumour, indicating clonal selection during recurrence in HCC. Overall, this approach provides a comprehensive and scalable solution to understand genome heterogeneity and evolution.

## Introduction

Heterogeneity is pervasive in human cancer [1] and manifests as morphologic, transcriptomic, and genetic differences between cells. However, intercellular genetic heterogeneity in cell populations is often obscured in genome analysis at the bulk level. Single-cell technologies have advanced rapidly in the past decade and can detect variants at the single-cell level [2–4]. Technologies for transcriptome analysis have been used to profile intra-tumour heterogeneity or define immune infiltration in various cancer types [5–13]. Although less widely utilized due to throughput and cost limitations, single-cell genome sequencing is a powerful tool to track clonal dynamics and infer evolutionary trajectories [14–18].

Most strategies for single-cell whole-genome sequencing (WGS) require whole-genome amplification (WGA) before library construction, which introduces bias and increases cost. The degenerate oligonucleotide-primed PCR (DOP-PCR) method attempts to amplify the entire single-cell genome by random oligonucleotide priming [19]. However, it preferentially amplifies regions rich in cytosine and guanosine, resulting in lower genomic coverage. Multiple displacement amplification (MDA) is another commonly used avenue utilizing random primers and the high fidelity φ29 polymerase. This method generates data with good genome coverage and lower error rates. However, due to the polymerase’s strand displacement activity [20], compromised uniformity is not suitable for copy number variation (CNV) detection. A hybrid method called multiple annealing and looping based amplification cycles (MALBAC) amplifies the genome with random primers and creates looped precursors to prevent continuous amplification before the PCR, achieving a better uniformity [21]. The other category of single-cell genome sequencing approaches is preamplification-free and transposase-based, including linear amplification via transposon insertion (LIANTI) [22], direct library preparation (DLP) [23], and transposon barcoded (TnBC) methods [24]. These approaches transpose single-cell genomic DNA directly and add common sequences to the end of the fragments for further amplification, reducing biases compared with preamplification-based techniques. These methods are based on a single tube or use complicated microvalve-based microfluidic chips, limiting the throughput.

Hepatocellular carcinoma (HCC) is a high-grade malignancy with a high recurrence rate of up to approximately 60% within 5 years [25]. As a risk factor for reduced survival, early recurrence of HCC is ascribed to a residual tumour and intrahepatic micrometastasis, closely related to intra-tumour heterogeneity [26]. Next-generation sequencing (NGS) studies based on cell population have reported a high degree of intra-tumour heterogeneity in HCC [27, 28]. A single-cell triple-omics approach applied to 26 tumour cells from HCC identified 2 tumour clones based on their CNV profiles [29]. Also, monoclonal and polyclonal origins have been reported recently based on single-cell WGS of ~ 30 cells in two individual patients [30]. However, a large number of cells are required to more comprehensively understand the heterogeneity in HCC, clonal expansion, and selection during HCC relapse.

Here, we developed an unbiased preamplification-free **s**ingle-**c**ell **D**NA library **p**reparation in **n**anolitre scale (scDPN) method using microwell chips and a 72 × 72 dual indexing strategy, which is capable of processing up to ~1,800 single cells in parallel. This approach can obtain highly sensitive and accurate single-cell CNV (scCNV) profiles based on evaluations in cell lines and tumour samples. We further applied this approach to paired primary and relapsed HCC tumour samples from the same patient. We identified 3 clonal subpopulations with aneuploid alterations across the genome. Furthermore, we noticed that relapsed tumour cells were originated from a minor subpopulation of the primary tumour, indicating clonal selection during HCC recurrence.

## Results

### Microwell-based single-cell DNA library preparation workflow

To increase scCNV detection efficiency, we developed a preamplification-free and unbiased single-cell DNA library preparation approach called scDPN for high-throughput scCNV detection, which provides a comprehensive, scalable solution for revealing genomic heterogeneity. The workflow of scDPN includes three main parts: cell isolation and single-cell identification, transposase-based (Tn5) library construction, and library pooling and sequencing. The first two steps were carried out in a 5,184 microwell chip (**Figure 1**). A cell suspension stained with Hoechst and propidium iodide (PI) was dispensed into the microwell chip with a MultiSample NanoDispenser (MSND). Cell suspensions with a range from 0.5 to 2.6 cells/50 nL (10~52 cells/μL) were optimum to obtain more than 1,000 wells with single-cell due to the cell counts per well followed a Poisson distribution. We used automated imaging to identify the number of cells and their viability, using fluorescent Hoechst and PI signals on a fluorescence microscope. Only microwells with single and viable cell (Hoechst^+^PI^-^) were selected for cell lysis and transposase fragmentation. Individual single-cell products were discriminated using 72 × 72 paired barcoded primers dispensed in succession with two individual dispensing steps. After several cycles of PCR, the barcodes and sequencing adaptors were added to both ends of the fragmented DNA. The microwell chip was then inverted, and all the barcoded libraries were collected into a pooled library. We determined the size distribution of pooled single-cell libraries by Agilent 2100 analysis (Figure S1). The libraries were then purified and cyclized for single-end 100 bp (SE100) sequencing on BGISEQ-500 [31].

**Figure 1.**
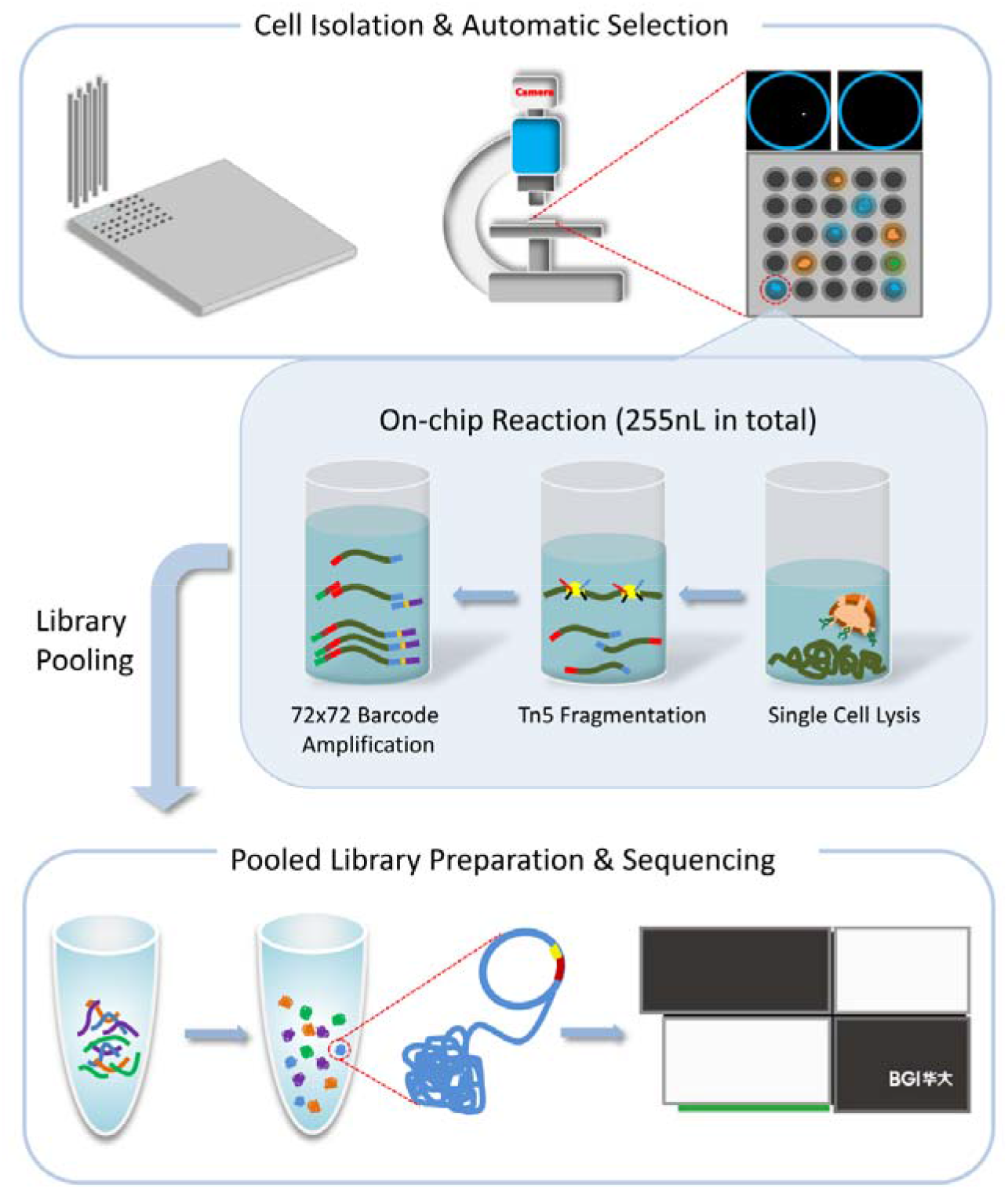
Schematic diagram of microwell-based single-cell genomic DNA library preparation. Stained cell suspensions were automatically dispensed into 72 × 72 microwell chips using MSND. Scanning fluorescence microscopy and cell selection software were used to discriminate wells containing single cells via the fluorescence of Hoechst and PI dyes. In the selected microwells, lysis buffer, Tn5 fragmentation buffer and 72 × 72 barcode primers were added step by step for single-cell DNA library amplification. The chip was incubated in a thermal cycler after each step. Indexed single-cell libraries were pooled by centrifugation for library purification, cyclized, and sequenced on the BGISEQ500 platform. PI, propidium iodide. MSND, MultiSample NanoDispenser.

### Assessment of data quality and uniformity under different reaction conditions

The HeLa S3 and YH cell lines, HCC adjacent normal liver tissue (ANT), and tumour tissues were processed and sequenced at 0.02× depth (~600k reads under SE100). To confirm whether our approach can generate enough data for scCNV detection, we draw a CNV saturation curve using three tumour cells with deeper sequencing depths up to 0.15× (**Figure 2A**, Materials and Methods). The number of detected CNVs increased in proportion to the number of randomly extracted uniquely mapped deduplicated reads (UMDR). When the amount of UMDR reached 300k, with an average sequencing depth of 0.01×, the detected CNV counts were saturated (Figure 2A).

**Figure 2.**
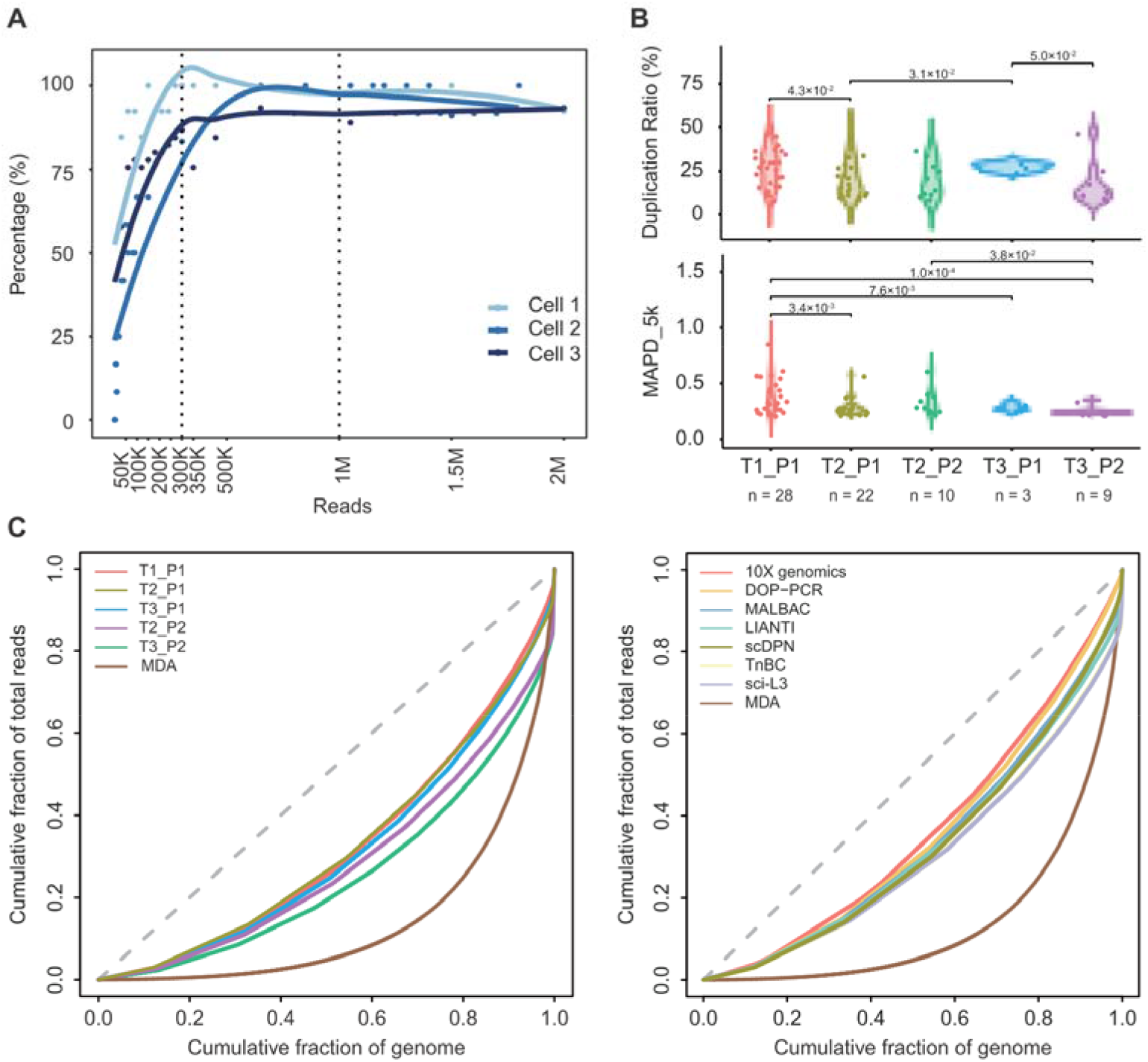
Assessment of library quality under different experimental conditions. **A.** CNV saturation curve. The detected CNVs increased with increasing numbers of unique mapped reads. At 300k reads, the CNV counts reach saturation. **B.** Sequencing data overview of 5 different single-cell lysis and transposase fragmentation conditions (T1_P1, n = 28; T2_P1, n = 22; T2_P2, n = 10; T3_P1, n = 3; T3_P2, n = 9). Violin charts showing the distribution of MAPD_5k and duplication ratio in different conditions with 400k raw reads. The Student’s T test was performed. **C**. Comparison of different library preparation conditions and the MDA method using Lorenz curves shows genome-wide coverage uniformity. The dotted straight black line indicates a perfectly uniform genome. **D**. Comparison of different library preparation methods (DOP-PCR, MALBAC, LIANTI, TnBC, sci-L3, and the 10x genomics) using Lorenz curves shows genome-wide coverage uniformity. The dotted straight black line indicates a perfectly uniform genome. CNV, copy number variation; MDA, multiple displacement amplification; DOP-PCR, degenerate oligonucleotide-primed PCR; MALBAC, multiple annealing, and looping-based amplification cycles; LIANTI, linear amplification via transposon insertion; TnBC, transposon barcoded; sci-L3, a single-cell sequencing method that combines combinatorial indexing and linear amplification.

We tested a combination of transposase (T1, T2, T3) and proteinase (P1, P2) reaction conditions to optimize the protocol. Single-cell libraries with raw data above 30k reads (5% of average reads) were assumed to have a template-based reaction, and 148 cells from 5 conditions were qualified (Table S1). Afterward, we selected the cells with oversaturated reads (UMDR > 300k) for further accuracy assessment. It was evident that condition T2_P1 (65%) showed the highest rate of cells passing the filtering criteria; conditions T1_P1, T2_P2, and T3_P2 showed a medium utilization rate between 40%~50%; and T3_P1 showed the lowest utilization rate, below 30% (Table S1). The qualified cells are listed in Table S2.

We statistically evaluated several features of these cells in different conditions, including mapped reads, coverage, duplication rates, and median absolute pairwise difference (MAPD) values. As the amount of sequencing reads affects these values, we down-sampled each single-cell library to 400k raw reads for comparison. Single-cell libraries treated with condition T3_P1 showed significantly fewer mapped reads and lower coverage (Figure S2A). A low duplication ratio reflects high data utilization. Conditions T2_P1, T2_P2, or T3_P2 had a mean duplication rate below 20%, which were lower than T1_P1 or T3_P1 (**Figure 2B**).

As a measurement of the bin-to-bin variation in read coverage, MAPD is an indicator of the evenness of WGA. Conditions T2_P1, T3_P1, and T3_P2 exhibited lower MAPD values (0.26 ± 0.07, 0.26 ± 0.03, and 0.23 ± 0.04, respectively, under 5k bins) compared with condition T1_P1 (0.37 ± 0.15 under 5k bins, *P*< 0.05) (Figure 2B). All of these conditions showed a much lower MAPD (mean MAPD < 0.4, 0.34 M mapping reads under a bin size of 300 kb) than that of normal cells prepared by MDA (MAPD: 0.4-0.6, 1.5 M mapped reads under a bin size of 500 kb [32]). We observed that CNV profiles generated from poor quality libraries had significant noise and larger MAPD values, so we set MAPD ≤ 0.45 as a cut-off for acceptable quality according to previous reports [32]. Because aberrant chromosomes influence MAPD, we compared the utilization rates from the same HCC tumour tissue under different conditions. The results showed that T2_P1 and T2_P2 had higher utilization rates up to 100%, by using a selection criterion of MAPD ≤ 0.45 for bin sizes of 600 kb or 300 kb (Figure S2B).

To further evaluate this approach’s genome-wide uniformity, we drew Lorenz curves for each condition and the data generated by the MDA method [24]. There were no substantial differences between the five conditions, and they all showed better uniformity than the MDA method (**Figure 2C**). Besides, the Lorenz curves demonstrated that scDPN had comparable uniformity with DOP-PCR, M ALB AC, LIANTI, TnBC, a single-cell sequencing method that combines combinatorial indexing and linear amplification (sci-L3) [33], and the 10x genomics CNV platform (**Figure 2D**). The T2_P1 condition was chosen as optimal for further applications.

### scDPN provides reliable data for accurate scCNV detection

To assess the sensitivity and accuracy of CNV calling with a depth of 300k reads, we first generated analogue data of CNVs of different sizes (1~15 Mb), with 20 variations generated for each size (Materials and Methods). Approximately 80% of CNVs above 2 Mb were detected in 5k, 10k, or 20k bins (Figure S3 A). The false discovery rate (FDR) was between 0.3~0.4 when detecting CNVs of 1 Mb and decreased to below 0.25 when detecting CNVs ≥ 2 Mb using 5k bins (Figure S3B).

To assess the approach’s reliability, we investigated the consistency of CNV profiles between single-cell and bulk populations. We used normal (YH) and tumour (HeLa S3) cell lines for single-cell copy number analysis and compared the results to the bulk CNVs from published HeLa S3 [34] and YH data [35]. HeLa S3 cells are known to harbour germline CNVs of defined sizes. The CNV profiles of single HeLa S3 cells were similar to the bulk data; however, this analysis did not detect a deletion on chromosome 4 posted in bulk HeLa S3 DNA (**Figure 3A** and S3C). We also observed different copy number states in chromosomes 13 and 18, which agreed with Liu’s discovery of substantial heterogeneity between HeLa variants from other laboratories [36]. The YH cells were B cells from a healthy donor, who was considered without significant CNVs. As expected, the single-cell YH cell CNV profile only had minor point CNV fluctuations (**Figure 3B** and S3C).

**Figure 3.**
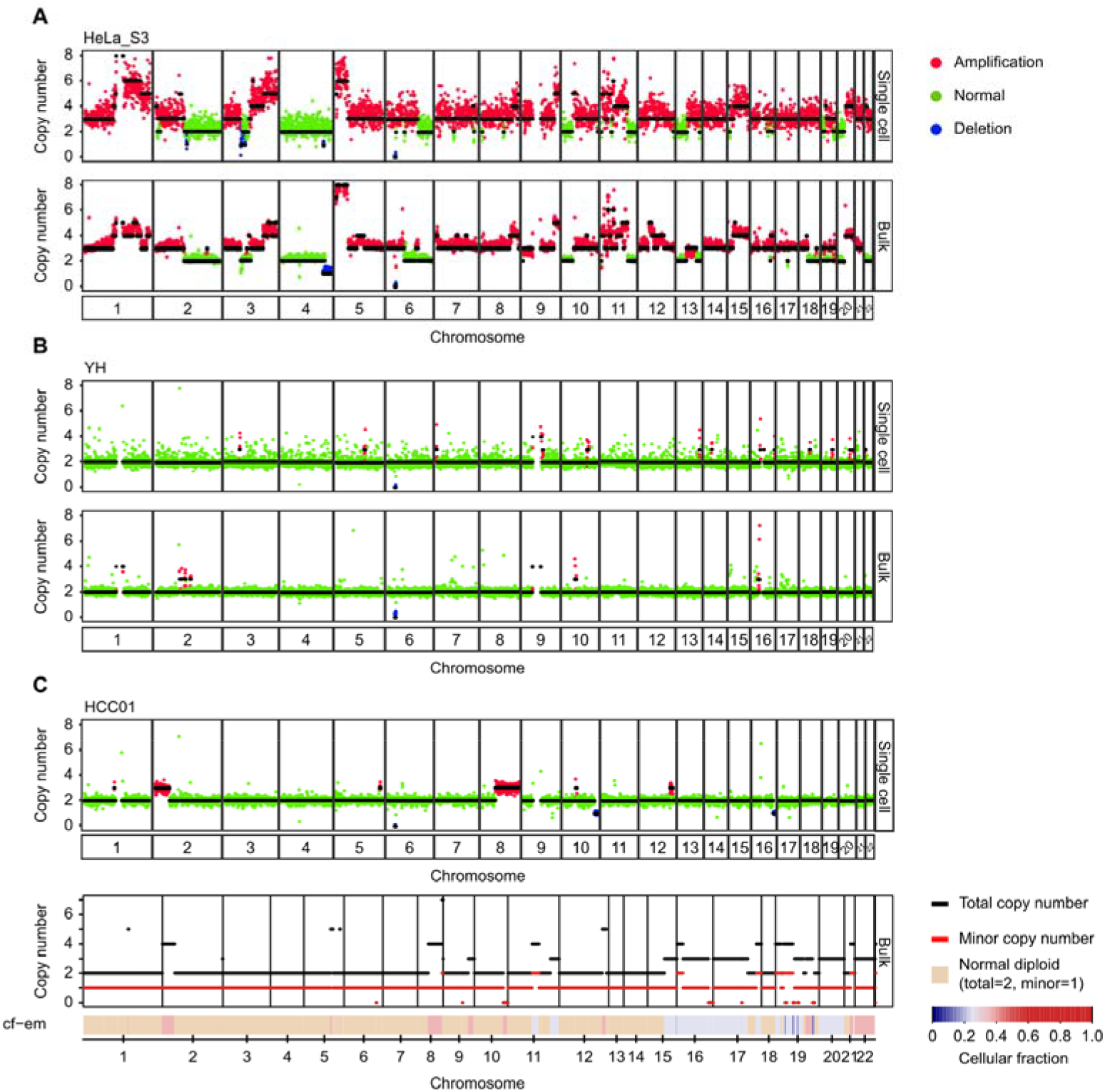
scDPN provides reliable data for accurate scCNV detection. **A.** Single-cell CNV profiles of HeLa S3 cells obtained using the T2_P1 condition and the corresponding bulk level HeLa S3 profile from published data. **B.** Single-cell resolution CNV profiles of the YH cell line obtained from the T2_P1 condition and the corresponding bulk level YH profile from published data. **C.** Representative single tumour cell copy number profile and corresponding bulk tumour CNV profile from FACETS analysis of whole-exome sequencing data in HCC01. The second panel plots the corresponding integer (total, minor) copy number calls. The estimated cellular fraction profile is plotted at the bottom, revealing both clonal and sub-clonal copy number events. HCC, hepatocellular carcinoma.

We then applied scDPN to an HCC tumour sample as well as paired ANT. The bulk tumour sample and peripheral blood mononuclear cells from the same patient (HCC01) were also subjected to whole-exome sequencing. We obtained 58 cells from HCC tumour tissue and 10 cells from ANT after filtering (> 300k reads, MAPD < 0.45). All 10 cells from ANT had no significant CNVs, as expected. One cell in the tumour did not have any CNVs and was considered normal (Figure S3C). The other 57 tumour cells had gain in 2p25.3-2p16.2, loss at 10q, and 56 had 8q11.23-8q24.3 gain (**Figure 3C** and S3C). This result indicated that there was only one major tumour clone in the HCC01 primary tumour. By comparing a representative copy number profile of a HCC tumour cell with a bulk CNV profile inferred from whole exome sequencing data (Materials and Methods), we observed concordant chromosome duplications of chromosomes 2, 8, and 12 and a deletion on chromosome 10, verifying the reliability of our CNV data. For example, the CNV profiles revealed multiple copy alterations, including 2p25.3-2p16.2, and 8q11.23-8q24.3, which are also present in the bulk DNA (Figure 3C).

### Single-cell CNV detection reveals tumour clonal subtypes in HCC

Genetic heterogeneity in HCC has been described in somatic nucleotide variations (SNVs) by NGS or SNP array of multiple regions from the same primary HCC bulk tumour tissue [37], but there are few studies at the single-cell level. Thus, we used scDPN to investigate tumour subclones in patient HCC02. After quality control (UMDR > 0.30 M, MAPD ≤ 0.45), we obtained 106 cells from the primary tumour for subsequent CNV calling. Three cells without chromosome copy number alterations were designated as normal cells. The remaining 103 cells showed two distinct CNV patterns, indicating that at least two tumour clones existed in this primary tumour (**Figure 4A**). The major subpopulation consisted of 87 cells with high-level amplifications on chromosomes 5p15.33-q35.3, 6p25.3-q12, 7p22.3-q36.3, 8q11.1-q24.3, and 15q11.2-q26.3 and deletions on chromosomes 6q12-q27 and 8p23.3-p11.21. Deletions of chromosomes 6q and 8p and gains in 6p and 8q are known recurrent CNVs in HCC [38]. A minor subpopulation of HCC02 comprised 16/103 (15.5%) tumour cells and had additional alterations: chromosome 1q21.1-q44 gain, 10q11.21-q23.31 loss, and 14q32.2-q32.33 loss (**Figure 4B**). We also observed common alterations in chromosomes 5, 6, 7, 8, and 15 in the same patient’s bulk tumour. However, the unique alterations in chromosomes 1, 10, and 14 found in the minor population of single cells were not detectable in the bulk tumour, demonstrating the capability of characterizing minor clones in single cells.

**Figure 4.**
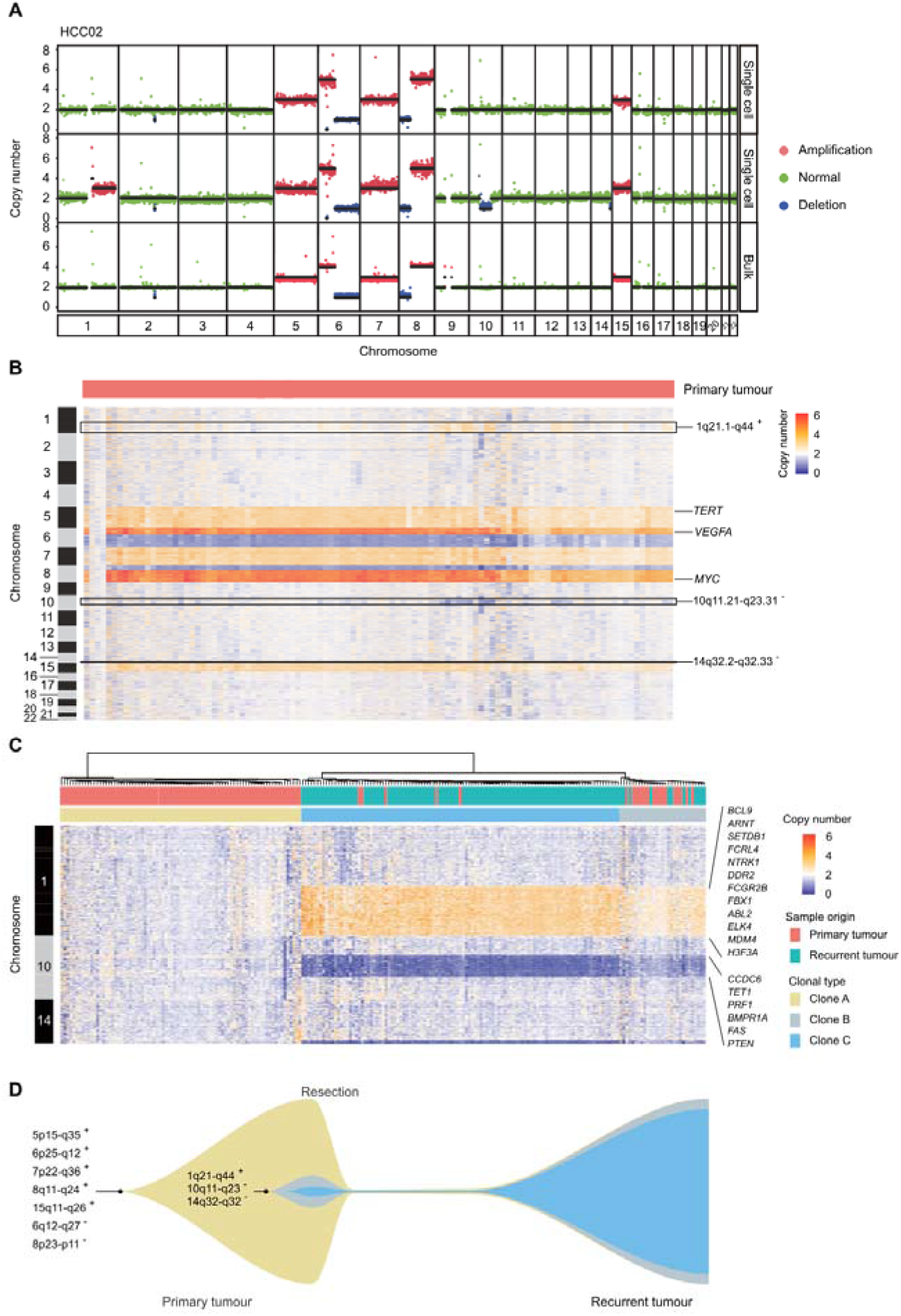
Single-cell CNV profiles reveal tumour clonal selection during HCC recurrence. **A.** Two CNV patterns observed in single-cell and CNV profiles detected by bulk WGS of the primary tumour in HCC02. Colours correspond to inferred copy-number states; black lines indicate segment medians. **B.** Heatmap showing the copy number states of all 106 cells from the primary tumour. Columns correspond to cells, and rows correspond to a ~600 kb genomic bin for each chromosome. Reported HCC-related genes *TERT*, *VEGFA*, and *MYC* are indicated. **C.** Heatmap showing the unsupervised clustering of all tumour cells from primary (n = 103) and relapsed tumours (n = 114) based on the CNVs on chr 1, 10, and 14. **D.** Schematic diagram of HCC tumour clonal selection during recurrence in patient HCC02.

### Clonal selection in HCC recurrence

A high recurrence rate is one of the risk factors contributing to the low 5-year survival rate in HCC. Understanding the clonal evolution and selection that occurs during relapse could aid in exploring the mechanism of recurrence. To investigate the correlation between the primary and recurrent tumour, we applied scDPN to the recurrent tumour from HCC02. We obtained 118 qualified cells from the recurrent tumour using the same filtering criteria. To our surprise, except for 4 normal cells without significant CNVs, the remaining 114 tumour cells had unique CNVs detected in the minor clone of the primary tumour, including 1q21.1-q44 gain, 10q11.21-q23.31, and 14q32.2-q32.33 loss (Figure S4A). This result strongly demonstrated that the minor clone in the primary tumour repopulated to be the dominant clone during relapse in this patient.

Furthermore, a hierarchical cluster analysis was conducted on CNVs in chromosomes 1, 10, and 14, revealing three subpopulations with distinct CNV patterns (**Figure 4C)**. Clone A comprised 81 primary tumour cells with no CNVs on these three chromosomes and corresponded to the major clone in the primary tumour. Both clones B and C showed similar CNVs in these three regions. Clone B was composed of 17 primary tumour cells and 12 recurrent tumour cells and was considered to be a transitional state of clone C. Clone C consisted of 102 relapsed tumour cells and 5 primary tumour cells, indicating that the minor clone in the primary tumour developed into a dominant clone during HCC relapse.

To determine which characteristics were associated with clone C selection during recurrence, we investigated the genes located in these unique CNV regions. We found several oncogenes and tumour suppressor genes described in the Catalogue Of Somatic Mutations In Cancer (COSMIC) database (Table S3). Several oncogenes were located in the amplification regions on chromosome 1q21.1-q44, including *ABL2, BCL9, DDR2, FCGR2B, ELK4,* and *MDM4,* while several tumour suppressor genes, including *PTEN, FAS,* and *PRF1,* were located in the loss region of 10q11.21-q23.31. We further validated that patients with 10q11.21-q23.31 loss or all the three alterations (1q21.1-q44 gain, 10q11.21-q23.31, and 14q32.2-q32.33 loss) showed lower disease or progression-free survival rate within two years in the TCGA dataset for HCC (Figure S4B). However, we did not observe a significant difference between patients with 1q21.1-q44 gain/14q32.2-q32.33 loss *vs.* others in disease-free survival, suggesting that the loss of 10q11.21-q23.31 may make a substantial contribution to tumour clone selection during relapse in HCC.

## Discussion

Single-cell genomic technologies have greatly aided the analysis of the evolution of cancer genomes and the study of genetic heterogeneity in cancer. However, the lack of high-throughput, cost-effective single-cell WGS approaches has limited their application. Here, we developed a preamplification-free, microwell-based single-cell DNA library preparation approach named scDPN, which can handle up to 1,800 cells per run. A fluorescence and imaging system enabled us to select a single and viable cell accounting for a lower doublet rate. Through a series of experiments, we determined the optimum on-chip experimental conditions for high data quality. The strategy for constructing libraries of scDPN was similar to the DLP and TnBC approaches. Improved version of LIANTI (sci-L3) and DLP (DLP+) [39] reported recently also have increased the throughput.

Compared with MDA methods, our platform generated single-cell genome data with better uniformity and lower noise, which decreases the required sequencing depth. Low-depth single-cell genome data of the HeLa S3 and YH cell lines and tumour samples generated by scDPN showed high sensitivity (only 0.02 × depth data needed) and accuracy compared with bulk tumour analysis. The small reaction volume substantially reduced the library construction costs to $0.5 per cell. ScDPN has the advantages of amplification uniformity, throughput, and cost over existing single-cell CNV detection methods. Additionally, we evaluated the performance of CNV detection in the cell nuclei from frozen tissues (Figure S5), which extends the application to additional cell types, including neurons and retrospective studies using frozen tissues.

However, scDPN is not suitable for SNVs detection due to low genome coverage. According to Zahn’s study, sequencing reads from all cells can be merged to produce a ‘pseudo-bulk’ genome with deep coverage accountable to an inference of SNV. Otherwise, a collection of high-depth ‘clonal genomes’ can be generated by combining all cells within a clone [23]. Additionally, there is a large difference in the amount of data among single-cell libraries produced from the same run due to the differential reaction efficiency during library preparation. Therefore, further condition optimization is essential to obtain uniform library products from an individual cell.

We used scDPN to identify subgroups of HCC tumour cells that were not detected in the bulk population (Figure 4A). This analysis indicates that important information is missing from bulk level-based sequencing studies. A large cohort based on scCNV in HCC may be needed to understand the genetic variance and heterogeneity more comprehensively. Understanding the clonal selection mechanisms in HCC recurrence could guide treatment and reduce relapse in HCC. Scaling our single-cell DNA preparation approach with paired primary and relapsed tumour samples could address essential questions concerning subclonal dynamics, such as how specific subclones evolve, evade immune surveillance, and metastasize.

In the profiling of CNVs in paired primary tumour cells (n = 103) and relapsed HCC tumour cells (n = 114), we observed a subpopulation (clone C) as the minor clone (5/103, 4.8%) in the primary tumour. This minor clone had additional CNVs of 1q21.1-q44 gain, 10q11.21-q23.31 loss, and 14q32.2-q32.33 loss, which developed into the dominant clone (102/114, 90%) in the recurrent tumour (Figure 4C). This a solid evidence to support the tumour clonal selection during HCC relapse (**Figure 4D)**. We validated in TCGA data that the loss of 10q11.21-q23.31, a region containing several tumour suppressor genes, is frequent in HCC and may play a crucial role in tumour clone selection during relapse. A chromosome 8p deletion has been correlated with HCC metastasis [40] and exists 3 clones in this tumour. The loss of 6p25.3-q12 presented in all clones would result in loss of heterozygosity (LOH) across the major histocompatibility complex (MHC), which is also reported to be associated with cancer metastasis [41]. Immune pressure has been proposed to shape the clonal evolution of metastasis [42]. However, the drivers or critical factors contributing to clonal selection during recurrence or metastasis in HCC and other cancers remain unclear. High-throughput single-cell omics from a large set of cancer patients, may potentially address these questions and simultaneously dissect the tumour environment, as well as the genetic and transcriptome characteristics of tumour cells.

## Materials and methods

### Cell line and patient tissue samples

The lymphoblastic cell line (YH cell line) was established from an Asian genome donor [35]. We purchased the HeLa S3 cell line from the American Type Culture Collection (CCL-2.2, ATCC, Manassas, VA, USA). The tumour sample used for on-chip reaction determination was a resected sample of a 45-year-old male patient (HCC01) with a primary HCC tumour. Paired primary and relapsed HCC tumour samples were obtained from a 63-year-old male patient (HCC02). Peripheral white blood cells and paired tumour sample and adjacent normal liver tissue were also obtained for bulk whole-exome sequencing or whole-genome sequencing.

### Preparation of the single-cell suspension

Cell suspension of cell lines were harvested and centrifuged at 500 g for 5 min, washed by phosphate buffer solution (PBS) buffer twice, and resuspended in PBS. The resected tumour samples were processed to a single-cell suspension using the commercial Tumour Dissociation Kit (30095929, Miltenyi Biotec, Bergisch Gladbach, Germany). Briefly, fresh tumour and adjacent normal liver tissues were cut into approximately 2-4 mm pieces and transferred into the gentleMACS C Tube containing the enzyme mix. Subsequently, the suspended cells were centrifuged at 300 g for 7 min after passing through cell strainers. The suspended cells were passed through cell strainers and centrifuged at 300 g for 7 min. The cell pellets were resuspended in 90% fetal bovine serum (FBS; 10270106, ThermoFisher Scientific, Waltham, MA, USA) with 10% dimethyl sulfoxide (DMSO; D8418-50ML, Sigma-Aldrich, St. Louis, MO, USA) and collected in a freezing container for −80 °C storage.

### Single-cell DNA library preparation and sequencing

We used the ReadyProbes Cell Viability Imaging Kit (R37609, ThermoFisher Scientific, Waltham, MA, USA) that contained Hoechst and PI to identify living cells. This staining process was at 37 °C for 20 min, then washed in cold 0.5× PBS twice. For cells from tumour tissue, we added fluorescent antibody CD45 (55548, BD Pharmingen™, San Jose, CA, USA) in the staining step. Based on FACS, CD45^-^Hoechst^+^ PI^-^ cells from the single-cell suspension were sorted into single tubes for tumour cell enrichment. Counted cells were dispensed into microwells using the ICELL8 MSND (640000, Takara Bio USA, Mountain View, CA, USA) at the concentration of 25 cells/μl in 0.5× PBS and 1× Second Diluent (640202, Takara Bio USA, Mountain View, CA, USA) into the ICELL8^®^ 350v Chip (640019, Takara Bio USA, Mountain View, CA, USA). We used the mixed buffer of PBS and fiducial mix (640202, Takara Bio USA, Mountain View, CA, USA) as the negative control wells. The MSND precisely dispensed 50 nL volumes into the microwells. Following cell dispensing, the chip was sealed with imaging film and centrifuged for 5 min at 500 g at 4 °C, and imaged with a 4× objective using Hoechst and PI. Following imaging, 35 nL cell lysis buffer was added to each microwell (P1: 2.89 AU/L Protease K (19155, Qiagen, Germany) and 72.8 mM pH 7.5 Tris•HCl (15567027, ThermoFisher Scientific, Waltham, MA USA); P2: 8.67 AU/L Protease K and 72.8 mM pH 7.5 Tris•HCl). The sealed chip was centrifuged for 3 min at 3,000 g and room temperature, then incubated at 50 °C for 1 h, followed by 75 °C for 20 min and finally 80 °C for 5 min to inactivate the protease. The chip was centrifuged for 3 min at 3,000 g again before 50 nL Tn5 transposition mix (T1: 0.06 U/μL Tn5 transposase (1000007867, MGI, China) and 2.4× TAG buffer (1000013442, MGI, China); T2: 2.4× TAG buffer, 0.14 U/μL Tn5 transposase; T3: 2.4× TAG buffer and 0.22 U/μL Tn5 transposase) were dispensed. After sealed, the chip was centrifuged at the same condition with the last step and incubated at 55 °C for 30 min. To stop transposase activity, 31 nL 5× NT buffer (0.25% SDS solution), 1.45 nL ddH_2_O, and 2.55 nL of Ad153-forward-tag (1~72) primer [1 μM] were dispensed, centrifuged and incubated for 5 min at room temperature. Another barcode primer was added to 50 nL PCR mix1 (29.6 nL 5× KAPA Fidelity Buffer, 7.69 nL 10 mM each dNTP, 5.1 nL PhoAd153 forward primer [10 μM], 5.1 nL Ad153 reverse primer [10 μM], and 2.55 nL of 72 Ad153-reverse-tag (1~72) primer [1 μM] made by KAPA HiFi HotStart PCR Kit (KK2500, Kapa Biosystems, Cape Town, South Africa). Finally, 50 nL PCR mix2 containing 21.4 nL 5× KAPA Fidelity Buffer, 5.1 nL 1 U/μL KAPA HiFi DNA polymerase, and 23.5 nL ddH2O was dispensed. We used the following conditions to perform PCR: 72 °C for 5 min; 95 °C for 3 min; 25 cycles of 98 °C for 3 min for 20 sec, 60 °C for 15 sec, and 72 °C for 25 sec; 72 °C for 5 min; and finally 4 °C. The final extraction of PCR products was carried out by centrifuging at 3,000 g for 3 min with an extraction kit. Product purification was performed using a 1.0× Agencourt Ampure XP bead (A63881, Beckman Coulter, Indianapolis, IN, USA) to sample ratio. Following ssDNA cyclization, digestion, and PEG32 bead purification (1000005259, MGI, China), the libraries were sequenced in SE100 on the BGISEQ-500 sequencer.

### Preprocessing of sequencing data

The raw reads derived from BGISEQ-500 were assessed by SOAPnuke (v1.5.6) [43] using the parameters “-Q 2 -G”. Afterward, we mapped the qualified reads to the human reference genome (hg19) by Burrows-Wheeler Aligner (BWA, v0.7.16a) [44] with BWA-MEM algorithms using arguments “-t 2 -k 32 -M /path/to/ref.fa”. The output SAM files were compressed and sorted by reference coordinates and then indexed with SAMtools (v1.1.19) [45]. Subsequently, uniquely mapped reads were extracted. Reads considered “PCR duplications” were removed by “samtools rmdup” from the downstream analysis.

### Detection of copy number variations

We calculated the copy number of each cell with an optimized method developed by the Baslan et.al. [35, 46, 47]. Based on the coverage suggestion of 30-180 reads per window for CNV calling from Gusnanto et al., we estimated the number of bins according to the average sequencing depth (< 1 Mbp) by the R package NGSoptwin [48]. The “bin boundaries” files for 5,000 bins in hg19 that suited the read length of 100 bp were generated. After GC content normalization, DNAcopy was employed for segmentation and copy number calculation, which points to gains and losses in chromosomes.

The FASTQ files of bulk HeLa S3 were downloaded from the NCBI Sequence Read Archive repository (SRP028541). The YH dataset was available in the GigaScience repository, GigaDB (http://gigadb.org/dataset/100115) [35].

For the matched bulk WES dataset, snp-pileup from htstools was first employed for processing BAM files using parameters “/path/to/dbsnp_150.common.hg38.vcf.gz -g -q15 -Q20 -P100 -r25,0”. We then used facets [49] for copy number estimation from the paired (normal/tumour) samples.

### Accuracy of CNV detection from the low coverage single-cell WGS data

The accuracies of CNV calling in the paper were assessed by sensitivity and FDR gain from the simulated dataset. A series of rearranged genomes with a defined size of CNVs was randomly generated by SimulateCNVs [50]. In each of the 10 outputs, 0.1 × WGS datasets with 20 CNVs of a specific size (1, 2, 3, 5, 10, 15 Mbp) were used to randomly extract 3 × 10^5^ uniquely mapped reads after duplicate removal with 5 replicates. A detected CNV was assumed to be true when it overlapped with at least 50% of the simulated CNVs. The sensitivity was defined as TP/(TP + FN), where the numerator was the true positive CNV mentioned above, while the total number of CNVs simulated served as the denominator. FDR was defined as FP/(FP+TP), where the numerator was the false positive CNV, and the denominator was the total number of CNVs detected by the algorithm.

### Estimation of sequencing saturations

The uniquely mapped reads after duplicate removal were randomly down-sampled to 3 × 10^4^, 6 × 10^4^, 9 × 10^4^, 1.2 × 10^5^, 1.5 × 10^5^, 1.8 × 10^5^, 2.1 × 10^5^, 2.4 × 10^5^, 2.7 × 10^5^, 3 × 10^5^, 4.5 × 10^5^, 6.5 × 10^5^, 1.05 × 10^6^, 1.5 × 10^6^, and 2 × 10^6^ reads. We used the down-sampled reads to estimate the sequencing saturation for our low-coverage WGS method. After calculating the copy number of each bin in the down-sampled datasets, the boundaries of the bins with copy number unequal to 2 were compared to that of samples with the highest read depth. The percentages of bins with abnormal copy number in samples with the highest coverage found in the down-sampled datasets were recorded. The saturation curves were fitted with locally weighted (LOESS) regression in geom_smooth function in the R package ggplot2 [51]. The inflection point of the curves was considered as the saturation point.

### Evaluation of the uniformity

The FASTQ files of MDA, DOP-PCR, MALBAC, LIANTI, TnBC, and sci-L3 datasets were downloaded from the NCBI Sequence Read Archive repository (SRR504711 for single-cell MDA, SRR1006146 for DOP-PCR, SRR975229 for MALBAC, SRX2660685 for LIANTI, SRX2847396 for TnBC, SRX5179905 for sci-L3) respectively. The sequence generated by 10x genomics platform was derived from https://support.10xgenomics.com/single-cell-dna/datasets/1.1.0/bi_cells_1k.

The uniquely mapped reads after duplicate removal from all samples were randomly down-sampled to 10^5^ reads for uniformity evaluation. To better indicate the bias of amplification methods, we binned reads into 60kb intervals across the genome with an average of 20 reads per bin according to the results from Xi et al. [24]. Reads in each bin were counted by bedtools2 (v2.20.1) and then applied for Lorenz model estimation.

### CNV profiling and tumor evolution visualization

MAPD is used for noise assessment in CNV calling [47, 52]. Since higher MAPD values reflect the poorer quality of a cell, we excluded single-cell samples with MAPD > 0.45. Segment ratios of samples were presented and clustered by hclust using ‘ward.D2’. Fishplot [53] was employed for fishplot construction.

## Ethical statements

We clarified that no animals were involved in this study. All samples involved in human beings were obtained after written informed consent and approval from the Institutional Review Board (IRB) at Fudan University ZhongShan Hospital and BGI-Shenzhen.

## Data availability

The low-coverage WGS data generated by BGISEQ-500 were deposited at CNGB Nucleotide Sequence Archive (https://db.cngb.org/) with the accession ID CNP0000448 and GSA at the National Genomics Data Centre (https://bigd.big.ac.cn/) with the accession ID HRA000478 and HRA000476.

## CRediT author statement

**Liang Wu**: Conceptualization, Methodology, Investigation, Writing-Original Draft, Writing-Review & Editing, Supervision. **Yuzhou Wang**: Methodology, Investigation, Writing-Original Draft. **Miaomiao Jiang**: Software, Data Curation, Investigation, Writing-Original Draft, Writing-Review & Editing, Visualization. **Biaofeng Zhou**: Software, Data Curation, Visualization. **Yunfan Sun**: Resources. **Kaiqian Zhou**: Resources. **Jiarui Xie**: Visualization. **Yu Zhong**: Software. **Zhikun Zhao:**Writing-Review & Editing. **Michael Dean**: Writing-Review & Editing. **Yong Hou**: Supervision, Project administration. **Shiping Liu**: Supervision, Project administration, Funding acquisition. All authors read and approved the final manuscript.

## Acknowledgements

This work was supported by Technology and Innovation Commission of Shenzhen Municipality (Grant No. GJHZ20180419190827179), and Science, Technology and Innovation Commission of Shenzhen Municipality (Grant No. JCYJ20170303151334808). We sincerely thank the support provided by China National GeneBank. We thank Dr. Xiaoyun Huang for the helpful comments on the manuscript. We are also grateful to Lei Li and Shishang Qin for assistance in data analysis as well as Dandan Chen for experimental support.

## Authors’ ORCID IDs

0000-0001-6259-261X (Liang Wu), 0000-0003-0033-6383 (Yuzhou Wang), 0000-0002-8473-8955 (Miaomiao Jiang), 0000-0002-0261-8256 (Biaofeng Zhou), 0000-0001-9790-2761 (Yunfan Sun), 0000-0002-2763-5976 (Kaiqian Zhou), 0000-0001-9022-4881 (Jiarui Xie), 0000-0001-6902-2732 (Yu Zhong), 0000-0001-5161-7818 (Zhikun Zhao), 0000-0003-2234-0631(Michael Dean), 0000-0002-0420-0726 (Yong Hou), 0000-0003-0019-619X (Shiping Liu).

## Supplementary materials

**Figure S1 Quality control of the library construction**

The length distribution of the library was determined using an Agilent 2100 bioanalyzer.

**Figure S2 Assessment for scDNP under different conditions**

**A**. Boxplots showing the distribution of mapped reads and genome coverage, in different conditions. The Student’s T test was performed. **B**. The proportions of HCC cells (UMDR > 300K) sampled from the same patient with MAPD </> 0.45 in different numbers of bins among various lysis and transposase fragmentation conditions (T1_P1, n = 22; T2_P1, n = 15; T2_P2, n = 5; T3_P1, n = 2, excluded; T3_P2, n = 4).

**Figure S3 scDPN provides reliable data for accurate scCNV detection**

Sensitivities (**A**) and FDRs (**B**) of the CNV detection algorithm at defined resolutions. The points and error bars represent the means and standard deviations, respectively. FDR, false discovery rate. **C**. Single-cell CNVs of different samples using low coverage. Heatmap showing the CNV profiling of HeLa S3 cells (red), YH cells (yellow), cells from adjacent liver tissue (blue), and tumour tissue (green) of HCC01. Columns correspond to cells, and rows correspond to 600 kb genomic bins for each chromosome. FDR, the false discovery rate.

**Figure S4 Tumour clonal selection during HCC recurrence**

**A**. Single-cell CNV profiling of HCC02 recurrent tumour samples. Heatmap showing the CNV profiles of all 118 cells from relapsed tumours. Columns correspond to cells, and rows correspond to 600 kb genomic bins for each chromosome. **B-C**. Kaplan-Meier analysis showing the disease/progression-free survival for patients with chr10q11.21-q23.31 deletion (**B**) and the three alterations (**C**) in the TCGA dataset for HCC.

**Figure S5 Evaluation of our CNV detection method with cell nuclei**

The distribution of mapped reads, used reads, genome coverage, and MAPD_5k of either the nucleus or cells are shown by box plots, and dots indicate individual samples. The Student’s T test was performed.

**Table S1 Statistics of cells used in the adjustment of reaction parameters**

**Table S2 Single-cell resource for scDPN assessment and tumour clone analyse**

**Table S3 Oncogenes and tumour suppressor genes with copy number alterations in our study**

